# Accurate measurement of microsatellite length by disrupting its tandem repeat structure

**DOI:** 10.1101/2021.12.09.471828

**Authors:** Dan Levy, Zihua Wang, Andrea B. Moffitt, Michael Wigler

**Affiliations:** Cold Spring Harbor Laboratory, Cold Spring Harbor, NY 11724, USA

**Author notes:** Authors contributed equally. Correspondence should be addressed to: Dan Levy.

## Abstract

Replication of tandem repeats of simple sequence motifs, also known as microsatellites, is error prone and variable lengths frequently occur during population expansions. Therefore, microsatellite length variations could serve as markers for cancer. However, accurate error-free quantitation of microsatellite lengths is difficult with current methods because of a high error rate during amplification and sequencing. We have solved this problem by using partial mutagenesis to disrupt enough of the repeat structure so that it can replicate faithfully, yet not so much that the flanking regions cannot be reliably identified. In this work we use bisulfite mutagenesis to convert a C to a U, later read as T. Compared to untreated templates, we achieve three orders of magnitude reduction in the error rate per round of replication. By requiring two independent first copies of an initial template, we reach error rates below one in a million. We discuss potential clinical applications of this method.

## INTRODUCTION

Tumors have genomic variants that distinguish them from the germline. These include single nucleotide variants (SNVs), small indels, large scale copy number variation (CNVs), and microsatellite length variation (MSLV). The profile of tumor variation has value in outcome prediction, the measurement of minimal residual disease, and possibly the early detection of cancer. In this paper, we focus on the accurate detection of MSLV. This class of variation is very abundant in the cancers of patients with defects in mismatch repair^1,2^, but also in cancers in general^3-6^. If MS lengths could be typed accurately it would open a potentially efficient way to fingerprint a cancer and to detect its presence in clinical specimens such as tissue biopsies, blood and urine^7,8^. The problem is that the same property of microsatellites that make their length unstable in cancer (and causes extensive heterogeneity in germline populations), namely, the repeat of a simple sequence motif, makes them unstable during amplification and sequencing. The microsatellite can expand or contract by units of the repeat, presumably due to polymerase slippage during replication^9-12^. This is particularly problematic when measuring the lengths of mononucleotide repeats, which are the most variable type of repeat in cancers^3-6^. In addition, modern day high-throughput sequence platforms fail to read through mononucleotide tracts accurately^13,14^.

To tackle this problem, various approaches have been tried. Multiplex PCR and capillary electrophoresis methods have been described to measure 5-10 microsatellite loci^15-17^. MS lengths have been characterized from gene panels and high throughput sequencing (HTS) data^7,18^, and statistical methods have been developed to increase accuracy in calling MS lengths from standard HTS data^19,20^. Droplet digital PCR has been employed^8^ to increase accuracy for small numbers of loci. None of these methods have the scale, depth, generality and accuracy needed for routinely monitoring a large panel of microsatellite loci deeply, and detecting minor variants.

Here we demonstrate a solution to the problem of accurate measurement of microsatellite length. We impose a partial random mutation pattern on templates^21^ and infer length from the reads containing the microsatellite only when the mutagenesis disrupts its repeat structure sufficiently to reduce error in amplification and sequencing. In the implementation described in the Results, we analyze three different microsatellites: mononucleotide tracts containing A or C, and a dinucleotide tract with a CA repeat. For the MS containing C, we randomly deaminate a proportion of the C’s to U’s, later read as a T, with a partial bisulfite reaction^22,23^. We add varietal tags (VT), random nucleotide sequences, to identify the initial templates and first copies of templates to measure and reduce sequence error^24,25,26^. We conclude with a discussion of the myriad potential applications of this method, and the obstacles that remain.

## MATERIALS AND METHODS

### Template design

For the testing and development of this method, we used three synthetic templates containing microsatellite (MS) tracts. The MS sequences are: a 17 base-pair mononucleotide A repeat called **M-17 (A)**, an 18 base-pair mononucleotide C repeat called **M-18 (C)**, and a 26 base-pair dinucleotide CA repeat called **D-26 (CA)**. The templates were ordered from Integrated DNA Technologies (IDT). The full sequences of the synthetic templates, oligonucleotide adaptors, and primers are given in **Supplementary Table 1**. As shown in **Figure 1**, the structure of synthetic templates used for the partial mutagenesis protocol, M-18 (C) and D-26 (CA), is as follows: a 5’ primer binding site without cytosine (UP1), a 15-mer varietal tag sequence (VT1) with random nucleotides represented as “NNN…”, a 5’ flanking sequence, a C or CA microsatellite tract, another 3’ flanking sequence, another 15-mer varietal tag (VT2) with random nucleotides represented as “DDD…”, and finally a 3’ binding site (UP2) without cytosine. We also examined templates containing mononucleotide A, denoted as M-17 (A), which did not undergo mutagenesis. These had a very similar design to the C microsatellite templates detailed in **Supplementary Table 1**. We use the notation of **M-18 (C+)** and **D-26 (CA+)** to denote the templates and libraries after mutagenesis, and **M-18 (C-), D-26 (CA-)**, and **M-17 (A-)** to refer to libraries without mutagenesis.

**Figure 1:**
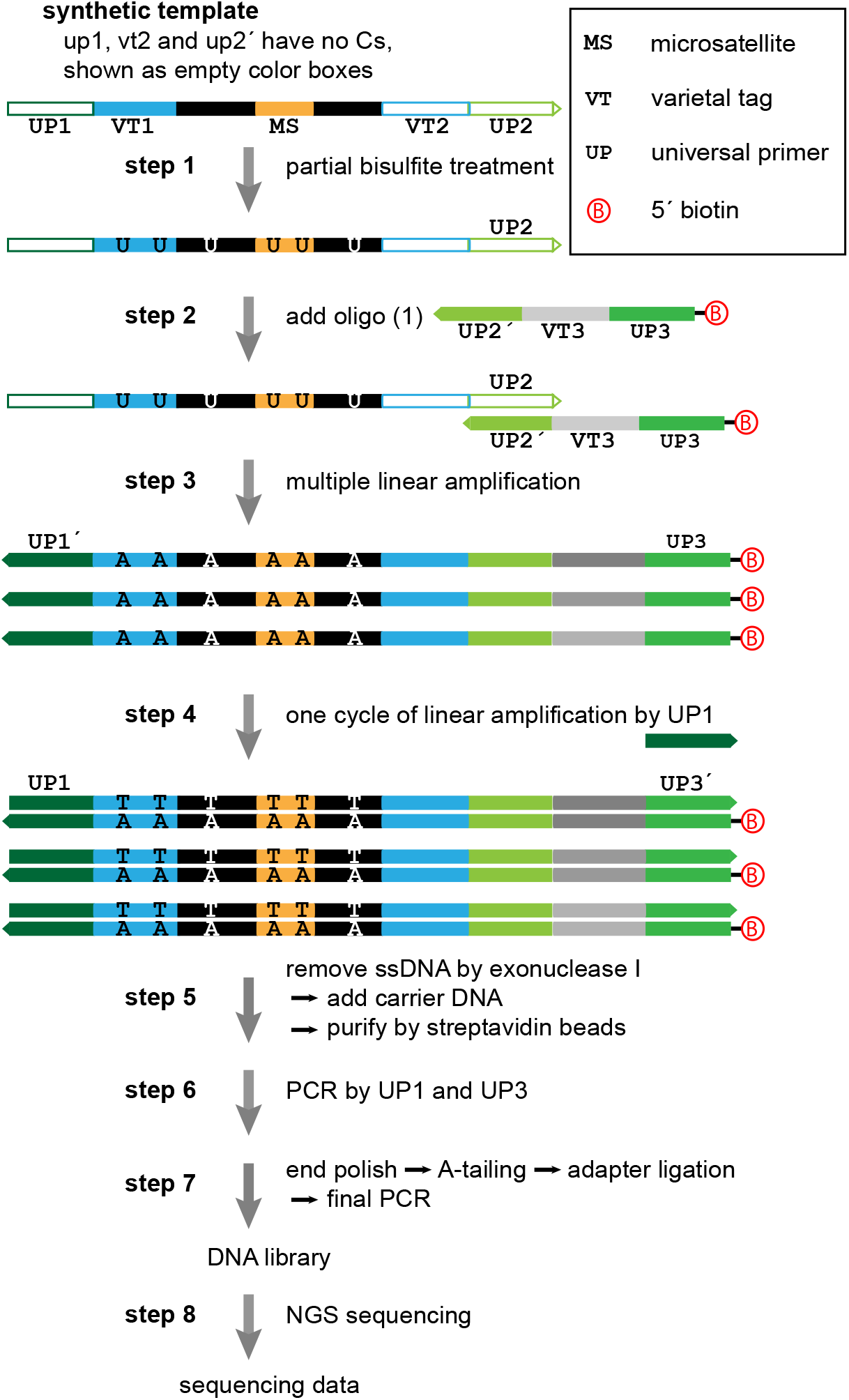
Bench protocol for partial mutagenesis. In step 1, each of the two synthetic templates containing C was partially bisulfite converted. In steps 2-3, about 6*10^4 of these templates underwent 9 cycles of linear amplification by using a biotinylated oligo. In step 4, double-stranded DNA fragments were obtained in another round of linear amplification by using UP1. In step 5, extra free oligo was removed by exonuclease I. After adding carrier DNA, biotinylated DNA fragments were purified by streptavidin beads. In step 6, the exponential PCR was carried out using UP1 and UP3 to generate enough material to prepare the DNA libraries for sequencing. These were sequenced as 2 × 150 bp paired-end runs on MiSeq (steps 7-8).

### Protocol for partial mutagenesis, library preparation, and sequencing

Our operational protocol for partial mutagenesis of microsatellite templates is described here and in **Figure 1**. In **step 1** of the mutagenesis protocol, 80 ng of each of the two templates containing C, M-18 (C) and D-26 (CA), was partially bisulfite converted (or not) by EZ DNA Methylation-Direct Kit (Zymo Research). Incubation time and temperature for bisulfite conversion were chosen to approach an ideal bisulfite conversion rate of close to 50%. In this protocol, DNA was incubated at 55°C for 40-50 min. In fact, we achieved 77% and 66% conversion for the M-18 (C+) tract and the D-26 (CA+) tract, respectively.

After conversion, about 6×10^4^ original templates underwent 9 cycles of linear amplification (**steps 2 and 3**) using a biotinylated oligo. This produced first round copies (**first copies**, for short) with a structure that had a 5’ biotinylated UP3 and a VT3 represented as “NNN…” in the **Supplementary Table 1** and in gray scale in Figure 1. Double-stranded DNA fragments were obtained in another round of linear amplification by using UP1 (**step 4**). In **step 5** extra free oligo was removed by Thermolabile Exonuclease I (NEB). After adding 50 ng of carrier DNA (poly (A), Sigma-Aldrich), biotinylated DNA fragments were purified by streptavidin beads (NEB). In **step 6**, 18 cycles of the exponential PCR were carried out using UP1 and UP3 to generate enough material to prepare the DNA libraries for sequencing. Standard steps for library preparation (end polishing, A-tailing, adapter ligation) were utilized to complete the sequencing library preparation (**step 7**). All libraries were prepared with variable length library barcodes^27^, and then pooled. The pooled libraries were sequenced as 2 × 150 bp paired-end runs on an Illumina MiSeq™ (**step 8**).

In steps 3 and 4 (linear amplification of first copies) we used NEBNext® Q5U® Master Mix. This master mix contains modified Q5® High Fidelity DNA Polymerase, optimized for amplification of uracil-containing templates. In step 6 we used Phusion Flash High-Fidelity PCR Master Mix (Thermo Fisher Scientific) for 18 cycles of PCR. This master mix contains Phusion Flash II DNA Polymerase which has high-fidelity and is excellent for multiplex PCR. In step 7, for library preparation, we used NEBNext® Ultra™ II Q5® Master Mix (NEB). This master mix contains Q5® High Fidelity DNA Polymerase, optimized for amplification of NGS libraries.

The parallel protocol without bisulfite treatment, used for the unmutated C templates, M-18 (C) and D-26 (CA) and M-17 (A), had the following differences: the number of original templates was about 3×10^4^, and in step 6, 14 or 15 cycles of PCR were employed.

### Sequence processing and tabulation

All read pairs were first evaluated for having the proper structure. A **proper** read pair has a good match (allowing up to one mismatch) to each of the UP1, UP2, and UP3 regions and the proximal flank of the microsatellite in both reads. From a proper read pair, we can extract the three varietal tags which identify the template (VT1, VT2) and first copy (VT3). From each read of the pair, we also searched for a good match to the distal flank sequence (up to one mismatch), and if the distal flank is found, we reported a microsatellite length (**MSL**) based on the distance in base pairs (bp) between the flanks within the read. A proper read always has a proximal flank, but it is possible that we could not clearly identify the distal flank. In those cases, the read did not report a length. We say a read pair is **qualified** if the both paired-end reads agree on the MSL, or if only one read of the pair reports a length. We then label these MSL as **on-target** if they are the expected length (i.e. 17, 18, or 26), and **off-target**, otherwise.

For qualified reads, we measured the degree to which the microsatellite is disrupted by the mutagenesis in two ways (**disruption indices**). The first is the **MS conversion rate** or the proportion of C bases converted to T in the MS, restricted to the tandem repeat. The second is the **maximum repeat length**, which is the largest number of tandem repeat units present in the disrupted microsatellite. We define a read as **sufficiently disrupted** if the MS conversion rate is between 0.15 and 0.85 **and** the maximum repeat length does not exceed five. See **Supplementary Table 2** for the expected yield of microsatellite disruption as a function of the average bisulfite conversion rate, as determined by simulation. Our observed levels of disruption follow closely the expectations from simulations.

We made a set of 5 tables from the proper reads for each of the 5 libraries (3 templates and 2 protocols): **M-17 (A-), M-18 (C-), D-26 (CA-), M-18 (C+), D-26 (CA+)**, which we call the READ TABLE (see Data Availability). The READ TABLE records the varietal tag information for the read, the MSL if the read is qualified (−1, otherwise), and its two disruption indices.

From the READ TABLES, we then made the FIRST COPY TABLES. A **first copy** is marked by its template tag-pair (VT1, VT2) and its first copy tag (VT3). We call a first copy **properly-covered** if it has a sufficient number of proper reads. The threshold for the number of reads required to define properly-covered depends on the complexity and depth of the library and was 10, 10, 20, 50, and 100 for M-17 (A-

), M-18 (C-), D-26 (CA-), M-18 (C+), D-26 (CA+), respectively. The choice for these cutoff, given the actual distribution of available reads, are shown in **Supplementary Figure 1**. We call a first-copy **well-covered** if it is properly-covered and has at least three qualified reads. For each well-covered first copy, we counted the MSL over all qualified reads to determine the **modal length**, which is the most common length reported by all reads associated with that first copy. A first copy is labeled **disrupted** if the median disruption parameters over its qualified reads are within the bounds to call a read sufficiently disrupted.

For each first copy, we tabulated the number of proper reads, the number of qualified reads, the modal length, and the number of qualified reads that report the modal length. For each first copy, we also recorded the median disruption indices of its qualified reads, where applicable. We then made the WELL-COVERED FIRST COPY TABLE by restricting to rows with a sufficient number of proper and qualified reads. This filtering step eliminated VT combinations with low read coverage that result from single-base errors in the varietal tag sequences.

From the WELL-COVERED FIRST COPY TABLE, we generated the TEMPLATE TABLE. A **template** is any template tag-pair (VT1, VT2) with at least one well-covered first copy. For each template, we counted the number of qualified reads, the number of well-covered first copies, and the median disruption indices from its first-copies. If those median disruption indices fall within the criteria defined for a sufficiently disrupted read, we call the template disrupted. We also record the modal lengths of the well-covered first copies for each template. We call a template **well-covered** if it has at least three well-covered first copies. Well-covered templates are flagged as **synthetic variants** if three or more first copies unanimously agree on an MSL length different from expected.

### Modeling error

We use two methods for modeling error. The first uses the method of **moments**; the second uses **Luria-Delbruck Diffusions, or LDD**^28^. The first method utilizes the varietal tags to identify reads with the first copies and the template from which they arose. Even if all templates are of the same initial length, the first copies may have different lengths due to replication error. Moreover, each first copy has its own probability of error due to random events occurring during exponential growth. We define the **N**-th moment for length **L** in the read data as the mean probability that N reads from each chosen first copy are unanimous for length L. Thus read error is nearly identical to the 1-st moment for any given L.

In the second method, we estimate per-round PCR error rates. In a (simple) LDD we model each round of an exponential amplification such that each single stranded nucleic acid of length **L** replicates with efficiency **e**, and its copy then retains its length, or decreases or increases its length by one unit according to two errors rate parameters: **α** and **β**. We assume for simplicity that α and β are not functions of strand or L, and the length increases or decreases only by the length of a single tandem repeat unit. All copies are retained in each round, but to simulate our protocol, the original template is not retained. After seeding with a single original template and **R** rounds of replication, we obtain a simulated distribution of MS lengths. We estimated α and β for our templates by simulating LDD distributions over a grid of parameter values and identifying the best match to the observed read error rate (see **Supplementary Data** for algorithm details).

## RESULTS

### Experimental Design

We evaluate the performance of the partial mutagenesis protocol for microsatellites by examining **5 sequencing libraries** generated from **3 synthetic templates** with and without bisulfite if the tract contained C. As shown in the top of **Figure 1**, the templates contain microsatellite tracts (mononucleotide A, mononucleotide C, or dinucleotide CA denoted **MS** and orange), flanked on either side by common sequence (black). 5’ and 3’ to the common sequence are varietal tag sequences (**VT1** and **VT2**, blue) that uniquely label each template molecule. Flanking the varietal tags are universal primer sequences (**UP1** and **UP2**′, green) designed without C nucleotides if needed to resist bisulfite mutagenesis.

Figure 1. shows our protocol for bisulfite mutagenesis, followed by library preparation and sequencing. The protocol is discussed in more detail in the Materials & Methods. Briefly, the templates were either bisulfite treated or not, in step 1. A biotinylated primer complementary to the 3′ universal primer UP2′, with its own unique varietal tag (**VT3**, gray) and a universal primer sequence (**UP3**) is added in step 2. We generate multiple **first copies** (step 3), each with the same **VT1-VT2** and a unique first copy tag **VT3**. The first copies are made double-stranded (step 4), purified by streptavidin chromatography (step 5), and amplified by PCR (step 6). Finally, the PCR products are made into libraries with specific barcodes, pooled and sequenced to high depth (steps 7-8).

The 5 sequencing libraries are named **M-17 (A-), M-18 (C-), D-26 (CA-), M-18 (C+), D-26 (CA+)**, corresponding to M (mono-) or D (di-nucleotide), their microsatellite length, the sequence of the microsatellite repeat unit, and whether mutagenesis was applied (+/-). These are abbreviated to **A-, C-, CA-, C+ and CA+**, respectively. A template is **disrupted** if the mutagenesis produces copies with a reduced tandem repeat structure (see Materials and Methods). In the analyses below, when we restrict the data for the mutated libraries to only the sufficiently disrupted templates, we refer to these datasets as **M-18 (C++)** or **C++**, and **D-26 (CA++)** or **CA++**. The proportion of sufficiently disrupted reads in the C++ and CA++ libraries is 29% and 73%, respectively, which falls close to the expected proportion, given our conversion rates (see **Supplementary Table 2**).

Below we describe the properties of the microsatellite lengths observed in the data for A-, C-, CA-, C++, and CA++, at three levels of organization: **(1) reads, (2) first copies, and (3) templates**. At the bottom level are the **templates** which refer to the original synthetized molecules. These are uniquely identified by their VT1-VT2 tag-pair. The next level is the **first copies** which are generated during the first round of linear amplification (Figure 1, step 2). First copies are identified by the unique triplet: the VT1-VT2 pair from their initial template, and the unique VT3 added to the molecule during linear amplification. At the top level are the **reads** of the sequencing library. For those reads with the correct structure, we determine its three varietal tags and, when possible, the length of the microsatellite. For details on data processing of reads, first copies and templates and for definitions of the terms used below, see Materials and Methods.

### Estimates of MSL Error Rates from Data

#### Reads only

Within a sequencing read, measuring the MSL depends on identifying the expected proximal and distal flank sequences and measuring their distance in the read. When parsing the structure of the unmutated mononucleotide reads, in particular the M-18 (C-) library, we found that the base quality of the read decays considerably after reading through the microsatellite sequence (**Supplementary Figure 2**). In many cases, this decay of base quality is so bad that the distal flank sequence could not be identified in the read. In the M-18 (C-) dataset, only 46% of proper reads are qualified, and almost all of these are from the read reporting G. In contrast, for the M-17 (A-) 95% of proper reads are qualified. For the remaining sets, 99% of reads are qualified (**Supplementary Figure 3**).

In **Figure 2, panels A-E**, we show the microsatellite length determinations per read as a histogram for each of the five libraries. In the histogram, qualified reads that match the expected length are shown in blue (**on-target**), while those reporting a different length are shown in orange (**off-target**). In general, off-target lengths tend to be shorter rather than longer. Read counts for each MS length and library are shown in **Table 1A**.

**Figure 2:**
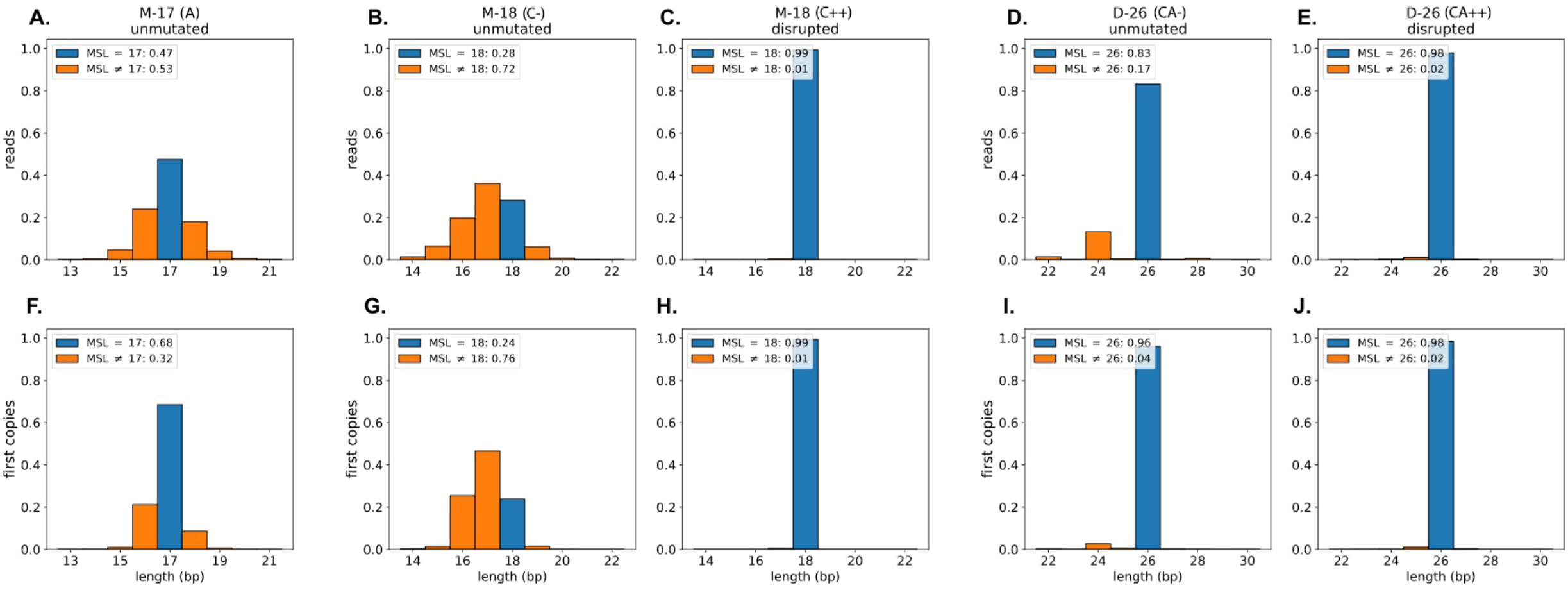
MSL distributions for reads and first copies. For each of the five libraries, we plot the distribution of observed MSLV. The upper panels show the distribution of read counts and the lower panels show the distribution of first copy modal lengths. The expected length is shown in blue, with off-target lengths shown in orange. The plot legends summarize the on and off-target rates per library. Data used to generate these plots is included in Table 1.

**Table 1:** MS length distributions in reads and first copies. For each of the five libraries, we show the number of reads and number of first copies that show each possible microsatellite length, from <= -5 to >= +5 from the expected length. For the three most stable libraries, where we were able to identify and remove synthetic variants, we show the resulting counts after removing synthetic variants.

| A. READS |  |  | MSLV deviation from expected length |  |  |  |  |  |  |  |  |  |  |
| --- | --- | --- | --- | --- | --- | --- | --- | --- | --- | --- | --- | --- | --- |
| | | | $\leq -5$ | -4 | -3 | -2 | -1 | 0 | +1 | +2 | +3 | +4 | $\geq +5$ |
| M-17 (A-) | unmutated | all | 1540 | 2382 | 15687 | 129032 | 662810 | 1318654 | 496859 | 114349 | 19831 | 3342 | 5390 |
| M-18 (C-) | unmutated | all | 4913 | 2511 | 10068 | 31175 | 56529 | 44531 | 9785 | 1360 | 229 | 51 | 200 |
| M-18 (C++) | disrupted | all | 1072 | 1 | 5 | 14 | 8451 | 1641719 | 60 | 2 | 0 | 0 | 0 |
| D-26 (CA-) | unmutated | all | 8957 | 37395 | 2357 | 339586 | 14455 | 2120035 | 5291 | 18719 | 18 | 702 | 121 |
| D-26 (CA++) | disrupted | all | 8395 | 206 | 848 | 2544 | 51425 | 4413601 | 14353 | 244 | 1 | 10 | 3 |
| M-18 (C++) | disrupted | wc - synth | 2 | 1 | 5 | 12 | 509 | 1612051 | 54 | 2 | 0 | 0 | 0 |
| D-26 (CA-) | unmutated | wc - synth | 3777 | 32367 | 413 | 303560 | 1321 | 1910332 | 450 | 16818 | 0 | 589 | 115 |
| D-26 (CA++) | disrupted | wc - synth | 61 | 196 | 2 | 2453 | 2076 | 4312537 | 405 | 236 | 0 | 10 | 3 |

| B. FIRST COPIES |  |  | MSLV deviation from expected length |  |  |  |  |  |  |  |  |  |  |
| --- | --- | --- | --- | --- | --- | --- | --- | --- | --- | --- | --- | --- | --- |
| | | | $\leq -5$ | -4 | -3 | -2 | -1 | 0 | +1 | +2 | +3 | +4 | $\geq +5$ |
| M-17 (A-) | unmutated | all | 96 | 58 | 180 | 2250 | 50762 | 164225 | 20588 | 1515 | 112 | 13 | 5 |
| M-18 (C-) | unmutated | all | 294 | 81 | 383 | 7171 | 13146 | 6717 | 428 | 11 | 0 | 0 | 4 |
| M-18 (C++) | disrupted | all | 7 | 0 | 0 | 0 | 58 | 11644 | 0 | 0 | 0 | 0 | 0 |
| D-26 (CA-) | unmutated | all | 86 | 82 | 23 | 1211 | 275 | 44142 | 95 | 36 | 0 | 2 | 1 |
| D-26 (CA++) | disrupted | all | 13 | 0 | 2 | 0 | 61 | 5852 | 18 | 0 | 0 | 0 | 0 |
| M-18 (C++) | disrupted | wc - synth | 0 | 0 | 0 | 0 | 0 | 11415 | 0 | 0 | 0 | 0 | 0 |
| D-26 (CA-) | unmutated | wc - synth | 6 | 54 | 5 | 1047 | 25 | 39557 | 8 | 32 | 0 | 1 | 1 |
| D-26 (CA++) | disrupted | wc - synth | 0 | 0 | 0 | 0 | 0 | 5680 | 0 | 0 | 0 | 0 | 0 |

For the M-17 (A-) tract, only 47% of the reads report the expected length of 17 bp. For the M-18 (C-) tract, the results are even worse: 28% of reads report the expected length. In contrast, the M-18 (C++) disrupted templates have 99% of reads on-target. For the D-26 (CA-) unmutated library, we find 83% of reads report the on-target length. The most frequently reported variants occur at 2 bp increments, equivalent to the size of the repeat unit. In contrast, the disrupted D-26 (CA++) templates have a high on-target rate with 98% reporting a length of 26. Unlike the unmutated D-26 (CA-), the off-target reads in D-26 (CA+) are almost entirely 1 bp off, reporting a length of 25 or 27.

#### First copies

We can, in principle, improve accuracy by taking a consensus of lengths from reads over first copies. We define the **modal length** of a first copy as the value most commonly seen among all the reads associated with it. The distributions of modal lengths are displayed in **Figure 2, in panels F-J**, and reported as counts in **Table 1B**. For the M-17 (A-) library, we see a slight improvement of the proportion of MS lengths on-target, from 47% when counting reads to 68% when counting the first copy consensus. For the M-18 (C-) library, we see a decline in lengths on-target from 28% to 24%. MS length estimates from the D-26 (CA-) unmutated library improve significantly when based on first copies, with 96% on-target, compared to 83% for reads alone. Unexpectedly, the lengths based on disrupted MS have nearly identical on-target rates, about 98% for D-26 (CA++) and 99% for M-18 (C++), whether using reads or first copy consensus. We suspected variation in the synthetic templates as the cause.

The proportion of **synthetic variants**, off-target length templates created during the synthesis of the original material, would not exceed the proportion of off-target first copies observed in the best data sets. This is the disrupted data: 2% for the D-26 (CA++) and 1% for the M-18 (C++). For the unmutated mononucleotide M-17 (A-) and M-18 (C-), the off-target rates are so high that they dwarf any potential improvement from removing these synthetic variant templates. However, identifying and removing the synthetic variant templates could dramatically improve the estimations of off-target rates of M-18 (C++), D-26 (CA++), and D-26 (CA-). To resolve this issue, we turn to the data aggregated over the initial templates.

#### Templates

For each initial template, we tabulate the modal lengths over all of its first copies. We condense this information by counting the number of first copies on-target (***x***) and the number of first copies that are off-target (***y***). **Figure 3** shows a scatter plot summarizing the distribution of (***x, y***) over all templates for each of the three libraries: M-18 (C++), D-26 (CA-), and D-26 (CA++). The size of the dot and intensity of the color reflect the proportion of templates with those values. For templates with no on-target first copies, we split the population between those whose first copies are <u>u</u>nanimous for an off-target length (orange, column U) and those whose first copies show <u>m</u>ultiple off-target lengths (column M). The underlying data are found in **Supplementary Table 3**.

**Figure 3:**
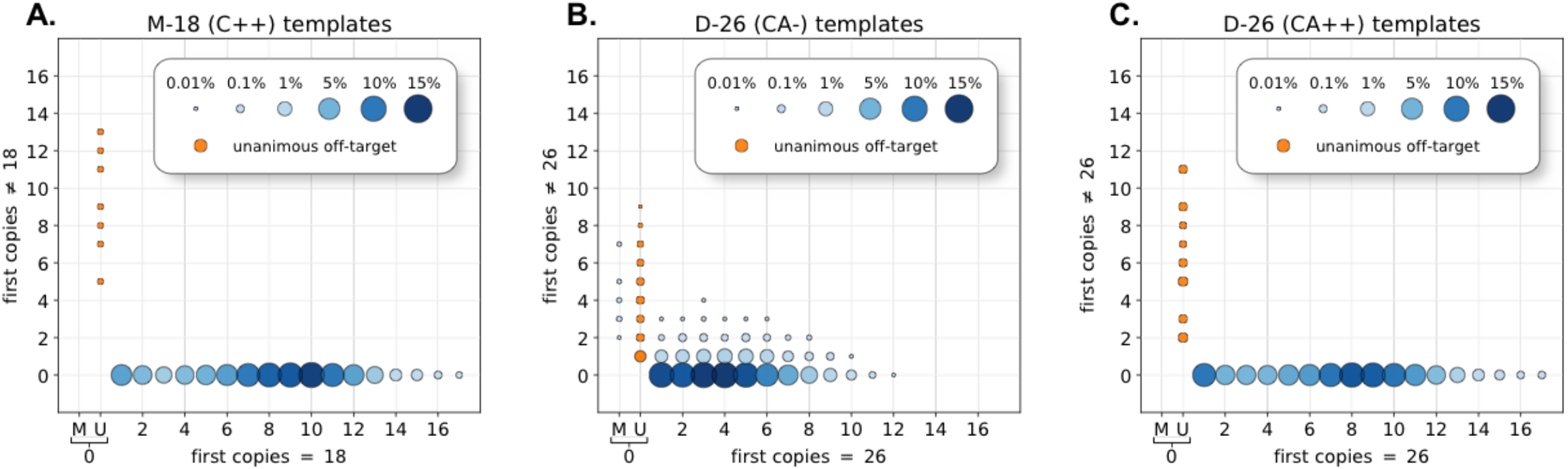
First copy consensus per template. For three libraries, we plot the distribution of templates with ***x*** on-target first copies and ***y*** off-target first copies. The size of the dot and intensity of the color reflect the proportion of templates, normalized by the total template count. Templates with no on-target first copies were further divided into “U” if all the first copies were unanimous for the same off-target length, or “M” if the first copies lengths were mixed. To highlight the population, unanimous off-target template populations are shown in orange. The underlying count data are available in Supplementary Table 3.

The templates with disrupted MS, both the mono-and di-nucleotide repeats, fall into two cleanly separable groups: the vast majority, in which the consensus MS length of first copies all unanimously agree with the on-target length; and a much smaller number of outlier templates, in which none of their first copies have an on-target length. All outlier disrupted templates were further examined. For each of these, all first copies unanimously agree on their unexpected MS length, most commonly one base-pair less than the expected length. The lengths of unanimous templates are shown in **Supplementary Table 4**.

The analysis of the unmutated D-26 (CA-) tract shows a more complex pattern (**Figure 3B**). As before, a majority of templates are on-target, with all first copies in agreement. There is also a small proportion of outlier templates, unanimous in that no first copies have the on-target length, and as before, most of these are unanimous for another length, typically one base less than the expected length (see **Supplementary Table 4**). It is noteworthy that relatively few of these outlier templates are +/-2bp the target length, as would be expected from polymerase error. In addition, there is a third and fairly numerous group of templates in D-26 (CA-): those with a predominance of first copies of the on-target length, yet containing one or more first copies with an off-target length.

#### Templates after removing synthetic length variants

Based on these studies, we labeled templates as synthetic variants if they had three or more first copies which were unanimous for an off-target length. For the three conditions just discussed, we removed these, and determined MS length error rates for the remainder. To do this fairly, we considered well-covered first copies and the reads associated with them only if from templates with at least three well-covered first copies. We estimate MS length read error from three of the five conditions shown in **Table 1A**, partitioning the error by its deviation from the expected length. The read error rates for disrupted reads in the M-18 (C++) and D-26 (CA++) data are on the order of 10^−3^ or better, but remain at about 16% for the undisrupted D-26 (CA-) data. Almost all of the erroneous reads in the D-26 (CA-) have MS lengths of 24. The erroneous reads for D-26 (CA++) are almost equally divided between lengths 24 and 25 (−2 and -1). The former value probably arises from residual tandem repeats following disruption. Consistent with this, we examined error rates as a function of disruption parameters (**Supplementary Table 5**), and note that if we had been even more restrictive, the read error rates could be reduced further.

In **Table 1B**, we similarly show first copy error, before and after removal for synthetic variants. After removal, we have 11,415 and 5,680 first copies in the M-18 (C++) and D-26 (CA++) data, respectively, 100% of which are of the expected length. First copy error for the CA-is reduced somewhat, but stands at about 3%. Most of the errors are to lengths of 24, two less than the target length, the error expected from slippage of one repeat unit.

### Limits of detection given error rates

So far, we have established that disruption reduces read error rates to 10^−3^ or better. In this section, we show that multiple disrupted reads, either from the same first copy or preferably from multiple first copies of the same initial template, achieve error rates on the order of 10^−6^ or better, whereas without disruption we do not better than ∼10^−2^ with CA-, the relatively stable repeat tract.

To measure error rates as a function of multiple reads we use the method of **moments**. For a given condition and template, we define the **N**^**th**^ **moment** of **L** as the probability of observing unanimous agreement of length L for N reads from the same first copy (see M&M). With moments, we can then estimate the probability of reads unanimous for length L with a configuration of **N[j]** reads over **J** first copies from the same template by multiplying the N[j] moments of L.

For the three conditions, CA-, CA++, and C++, we show the moments in **Table 2** for L equal to the expected lengths (26, 26 and 18) and for L the most commonly observed length in error, a one-unit deletion (24, 24 and 17, respectively). Note that the N-th moments are not powers of the first moment, because the moments account for variation in error rates across first copies. This variation is inherent in exponential growth, a mathematical insight first noted by Luria and Delbruck^28^.

**Table 2:** Luria-Delbruck Diffusion simulation parameters and results. For the M-18 (C++), D-26 (CA-), and D-26 (CA++) conditions, the table shows the Luria-Delbruck Diffusion (LDD) per-round error rate parameter selected to best fit the observed data. (Left) The results of the simulation are shown as number of reads and first copies at length 26 and 24 (or 18 and 17), for each condition. (Right) Parallel results after downsampling from observed data. The simulation total counts are matched to the downsampled total counts. Further, the table shows the proportion of first copies at each length, given N number of unanimous reads for both simulated and downsampled data. Off-target counts in the disrupted libraries are 0, due to the extremely low error rates of these conditions, necessitating the estimation of error via simulation to accurately estimate error rates.

| **A.** |  | **D-26 (CA-)** |  |  |  |
| --- | --- | --- | --- | --- | --- |
|  |  | **observed** |  | **LDD simulated, $\alpha = 1.35\text{E-}02$** |  |
| **MS length (L)** |  | **24** | **26** | **24** | **26** |
| **number of reads** |  | 303,560.00 | 1,910,332.00 | 308,265.08 | 1,920,815.70 |
| **number of first copies** |  | 1,047.00 | 39,557.00 | 957.93 | 39,733.44 |
| **N-th moment**<br>**for length L** | **N = 1** | 0.133978 | 0.841325 | 0.135544 | 0.846076 |
|  | **N = 2** | 0.033846 | 0.729053 | 0.029580 | 0.731265 |
|  | **N = 3** | 0.017401 | 0.636207 | 0.013170 | 0.635169 |
|  | **N = 4** | 0.013061 | 0.558259 | 0.009098 | 0.553591 |

| **B.** |  | **D-26 (CA++)** |  |  |  |
| --- | --- | --- | --- | --- | --- |
|  |  | **observed** |  | **LDD simulated, $\alpha = 4.95\text{E-}05$** |  |
| **MS length (L)** |  | **24** | **26** | **24** | **26** |
| **number of reads** |  | 2,453.00 | 4,312,537.00 | 2,482.78 | 4,315,256.48 |
| **number of first copies** |  | 0.00 | 5,680.00 | 0.41 | 5,679.54 |
| **N-th moment**<br>**for length L** | **N = 1** | 0.000578 | 0.998725 | 0.000575 | 0.999370 |
|  | **N = 2** | 0.000005 | 0.997459 | 0.000072 | 0.998820 |
|  | **N = 3** | 0.000000 | 0.996201 | 0.000055 | 0.998289 |
|  | **N = 4** | 0.000000 | 0.994952 | 0.000050 | 0.997771 |

| **C.** |  | **M-18 (C++)** |  |  |  |
| --- | --- | --- | --- | --- | --- |
|  |  | **observed** |  | **LDD simulated, $\alpha = 2.80\text{E-}05$** |  |
| **MS length (L)** |  | **17** | **18** | **17** | **18** |
| **number of reads** |  | 509.00 | 1,612,051.00 | 532.80 | 1,612,048.90 |
| **number of first copies** |  | 0.00 | 11,415.00 | 0.52 | 11,414.43 |
| **N-th moment**<br>**for length L** | **N = 1** | 0.000324 | 0.999631 | 0.000330 | 0.999637 |
|  | **N = 2** | 0.000005 | 0.999268 | 0.000045 | 0.999323 |
|  | **N = 3** | 0.000000 | 0.998913 | 0.000035 | 0.999020 |
|  | **N = 4** | 0.000000 | 0.998563 | 0.000032 | 0.998724 |

In the CA-data, the first moment for 24 matches the average read error rate of 1.3×10^−1^. The second moment for that length in CA-diminishes to 3.4×10^−2^ reflecting increased accuracy from two unanimous reads. The third and fourth moments continue the downward trend but will plateau, reflecting that for large N, the moments cannot be lower than the first-round error rate. We estimate this from the data to be about 1.3×10^−2^ for CA-.

For the disrupted datasets, CA++ and C++, the first moments for off-target also match the average read error rates of 5.8×10^−4^ and 3.2×10^−4^, respectively. The probability of seeing two unanimous off-target reads, the 2-nd moment, dramatically decreases to 5×10^−6^. The higher moments are vanishingly small. However, due to the number of observable first copies (5680 for CA++ and 11415 for C++, see Table 1B) and low error rate, we see no first-round error in any copy (Figure 3), so these values cannot plateau. We felt it likely therefore that we were underestimating the tail of the distribution that define the higher moments.

To obtain a better approximation for the higher moments, we use a Luria Delbruck Diffusion (LDD) model that simulates error during amplification (see M&M for details). In addition to providing a simulation of the higher moments, a good LDD fit to the data can provide an estimate of per round error rate. With a good estimate of the per round error rate, we can also accurately simulate any number of rounds of amplification.

A simple LDD model has four parameters: an efficiency of replication (**e** = 0.95), the number of rounds of replication (**R** = 23), and two **per round length error rates**: one unit decrease (**α**), and one unit increase (**β**). For each dataset, we numerically generate LDD distributions over a grid of α and β, and select the grid-point where the read error-rates (1^st^ moments) best match the empirical data. **Table 2** shows the LDD simulation data for the best-fit α parameter for each dataset: the first copy counts on-and off-target by one unit for a given read count (top two rows) and the four moments, on and off-target by one unit.

Comparing observed to LDD simulation in Table 2, we find that the expected number of reads agree closely, as do the first moments. This is unsurprising because we choose the LDD parameters to fit these values. However, the LDD simulations provide good estimates for the expected number of off-target first copies and the higher moments of the CA-dataset. The zero observations of off-target first copies in the disrupted CA++ and C++ data, which resulted in underestimated higher moments, are approximated in the LDD simulations by 0.41 and 0.52 respectively. This results in a more reasonable numerical behavior of the higher moments, now bounded by a nonzero first round error rate (**α**).

Using the LDD sample moments, we can now estimate detection limits for various unanimous read configurations. Six examples are shown in **Table 3** for CA- and CA++ templates. It is clear that the disrupted microsatellites are measured with at least three orders of magnitude lower error than undisrupted microsatellites. Moreover, obtaining reads from the same first copy has a higher error rate than the same number of reads from multiple first copies. Given three unanimous reads over two first copies, less than one in ten million CA++ templates with MSL of 26 would be mistaken as having a MSL of 24. Under conditions of disruption, we estimate a limit of detection at or below one in a million, enabling highly sensitive detection of rare microsatellite length variants in biological samples.

**Table 3:** Error rates with unanimous lengths by condition and read configuration. For the D-26 (CA-) and D-26 (CA++) conditions, we use the LDD simulation results to show expected rates of observing length 24 in error, depending on the number of first copies and the read configuration across those first copies. Error rates are orders of magnitude lower when reads are distributed over multiple first copies.

|  |  |  | D-26 (CA-) |  | D-26 (CA++) |  |
| --- | --- | --- | --- | --- | --- | --- |
|  |  |  | unmutated |  | disrupted |  |
| configuration | reads | first copies | 24 | 26 | 24 | 26 |
| 1 | 1 | 1 | 1.4E-01 | 0.846 | 5.7E-04 | 0.999 |
| 2 | 2 | 1 | 3.0E-02 | 0.731 | 7.2E-05 | 0.999 |
| 3 | 3 | 1 | 1.3E-02 | 0.635 | 5.5E-05 | 0.998 |
| 1, 1 | 2 | 2 | 1.8E-02 | 0.716 | 3.3E-07 | 0.999 |
| 2, 1 | 3 | 2 | 4.0E-03 | 0.619 | 4.2E-08 | 0.998 |
| 1, 1, 1 | 3 | 3 | 2.5E-03 | 0.606 | 1.9E-10 | 0.998 |

### Determining error from paired first copy observations

In the above sections, we established error rates of reads per template and per first copy by analysis of deep coverage data (**Table 1**). We determined the presence of synthetic variants, and determined their lengths and proportion (**Supplementary Table 4**). We demonstrate that when reads from at least two distinct first copies are in unanimous agreement, the false positive rate is many orders of magnitude lower when the simple repeat structure of a microsatellite is disrupted than when it is not (Table 3). This analysis indicated that single read coverage over two to three first copies for individual templates in a panel of assayed loci will suffice to give accurate measure of rare variant frequency. We show in the following that we can indeed measure error rates and confidently detect rare variants from **paired first copy observations**: one read over each of two first copies, by emulating low coverage from our data set without removal of synthetic variants.

We counted all paired first copy observations **(L, K)** where **L** is the MS length from one read, **K** the length from the other. To create an emulation in which all templates are weighted equally from the entire data set, we normalized counts per template, that is, for every template we determined the number of paired counts that were **(L, K)** and divided by the total. This generates a matrix over all possible **(L, K)** pairs for every template. These distributions are then averaged (**Supplementary Table 6**). To emulate low-coverage paired first copy observations over N templates, we multiply this table by N, and round to the nearest integer. We did this for each of the unmutated and disrupted conditions (C-, C++, CA-, and CA++) for N = 10^4^ templates (**Supplementary Table 7**).

We illustrate the method of error estimation for the pairs (L = 26, K = 24) from the D-26 template, both unmutated (CA-) and disrupted (CA++), and for the pairs (L = 18, K = 17) from the M-18 template disrupted data (C++) shown in **Table 4**. As described above, we emulated 10,000 templates, each with two reads from different first copies, and show the counts for the relevant (L, K) pairs. When the counts of (L, L) >> (K, K), the proportion of (L, K) is, to a first approximation, 2\**p*\*(1-*p*) where *p* is the proportion of reads that report K rather than L. This equation depends upon the assumptions that all K derive from L; that reads drawn from different first copies have independent error; and that we can ignore the proportion of L that derive from K. If *x* is the proportion of (L, K) pairs, then 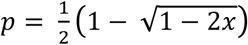.

**Table 4:** Error rates from paired first copy observations. For the M-18 (C++), D-26 (CA-), and D-26 (CA++) conditions, we generated an emulated dataset consisting of 10,000 pairs of reads from independent first copies. MS lengths are compared between reads in a pair to determine read error and the expected number of templates that agree at the unexpected length K. Shown are the number of emulated templates with lengths L, L and lengths L, K; the read error from paired first copy observations compared to the LDD moment method; the number of K,K templates expected by error, from synthetic error, and in the emulated dataset. The last row shows the likelihood of observing zero K,K pairs by error alone.

|  | **D-26 (CA-)**<br>L=26, K=24 | **D-26 (CA++)**<br>L=26, K=24 | **M-18 (C++)**<br>L=18, K=17 |
| --- | --- | --- | --- |
| emulated (L, L) | 6999 | 9766 | 9943 |
| emulated (L, K) | 2240 | 11 | 6 |
| read error $p$ | 0.13781 | 0.00056 | 0.00030 |
| first moment (Table 2) | 0.13712 | 0.00057 | 0.00032 |
| (K, K) estimate from $p$ | 179 | 0 | 0 |
| (K, K) estimate from synth (Supp. Table 4) | 0 | 0 | 43 |
| emulated (K, K) | 187 | 0 | 43 |
| likelihood (K, K) = 0 from $p$ | 0.00000 | 0.99691 | 0.99910 |

As seen in **Table 4**, the derived *p* error rate from low-coverage pair data agree with the meticulously derived read-error rate (first moments) from **Table 2**, which involved removing synthetic variants. For the unmutated CA-data, the 14% read-error rate predicts that 24-24 paired first copy observations should arise from length 26 templates at a rate of 1.9%. Given binomial sampling from the 10,000 templates, we expect that 179 K-K pairs in our emulation arise from error, with a 90% confidence interval of +/- 20 pairs. In the emulation, we observe 187 K-K pairs, well within the bounds explained by measurement error. We note that small deviations from the expected count would be indistinguishable from noise.

In contrast, the read error rates for the disrupted CA++ microsatellites are less than 10^−3^, so that the chance of two independent errors are below 10^−6^. For disrupted templates, the expected count of 24-24 pairs in our emulation of 10,000 templates is a firm zero, with the probability of observing 0 such pairs from error of 0.997. From the high-coverage data over unanimous templates (**Supplementary Table 4**), we do not see any deviations from this count. In contrast to the unmutated data, a count of 1 would represent confident detection.

Similarly, in the mononucleotide C++ library, the reads of the disrupted template (with MSL 18) are rarely read as 17. The estimated read-error rate is below 10^−3^ and our expectation of observing 17-17 from length 18 templates is 0 with very high confidence (0.9991). In our emulation of 10,000 templates, we do observe 17-17 pairs, 43 in total. Given the low probability of this occurring by error, all 43 of those templates (0.43%) are likely real and represent variant synthetic templates with MSL 17. As confirmation, this number precisely matches our expectation of synthetic variants based on high-coverage unanimous templates (6 / 1397 = 0.429%, Supplementary Table 4).

## DISCUSSION

The length of a microsatellite is unstable during replication. On the one hand, the high rate of variability makes microsatellite lengths attractive markers for disease; on the other hand, instability makes accurate microsatellite length measurement very difficult^9-12^. This is especially true for the mononucleotide repeats, the most highly variable microsatellites and potentially the most valuable markers. The measurement of mononucleotide repeat length is further aggravated by modern high-throughput sequencing platforms^13,14,29^. As we discuss in the Introduction, various attempts have been made to ameliorate some of these problems^8,15-17^. However, none of these approaches have the breadth, depth and sensitivity to detect minor populations of variant lengths over many thousands of templates and loci in a single assay.

We solved this problem by conceiving and implementing the strategy of measuring microsatellite length in templates by first disrupting the very structure which renders their replication unfaithful. We disrupted the tandem repeats in the templates using bisulfite conversion of C to a U, later read as a T. Partial mutagenesis is critical, as when the mutagenesis is too complete, a new repeat structure is created, and replication again becomes unreliable. Thus, our method depends on determining the degree of disruption of the repeat structure in each sequence read. We developed two disruption indices, and used them both to identify templates with lower error rates, but we are still exploring other methods for quantifying the disruption of a repeat structure.

To demonstrate the ability of partial mutagenesis to stabilize microsatellites for amplification and sequencing, we created a controlled test system using synthetic templates. Each template was synthesized with an identifying tag, and we made multiple independent first copies, each with its own identifying tag. We determined the fidelity of replication as a function of the degree of disruption of the repeat structure. By aggregating over the individual first copies and the individual initial templates with extremely high depth of coverage, we were able to remove templates that were synthetic variants, and then to determine with great precision the error rate in microsatellite length measurement. Fitting observations to what we call Luria-Delbruck distributions, we estimated the length error per round of replication.

We obtain two to three orders of magnitude improvement in per round error when the repeat structure is disrupted compared to when it is not, with the greatest improvement seen in mono-C tracts. Without disruption, mono-C tracts have very high error rates, and the sequencers often fail to get a robust sequence from a read. Under the conditions of our assay, we predict that error rates will be less than 10^−6^ with just a few reads distributed over a few independent first copies. We confirmed these predictions on an emulated data set of low coverage per template, in which synthetic variants were not removed. Using a method based on two reads per template from independent first copies (i.e., paired first copy observations) we confirmed our estimated error rates and correctly identified the proportion of synthetic variants, analogous to the detection of biologically derived rare variants in a sample.

The value of accurate MSLV determination stems largely from its many applications in cancer^3-6^. In particular we are interested in two goals: measuring the load of a cancer with known genomic variation; and detecting cancer early in persons at risk. The first goal has been achieved using patient specific single nucleotide variations (SNVs)^26,30-34^. But while SNVs are sparse and can occur anywhere in the genome, MSLV are frequent and occur at known loci. Therefore, given enrichment for those loci with panels and a reliable method for measuring length, an assay based on microsatellites would be much less expensive than the alternatives, standard for every patient, and could be more rapidly deployed. Moreover, an assay based on a panel of many variable loci might well detect the emergence of a new variant that potentially indicates the escape of a new clone from therapeutic control.

A transformative value for MSLV detection may lie in its application to early detection. Many studies, including our own unpublished work, indicate that upon presentation with disease most if not all patients have traces of their cancer genome in the cell free component of blood. It follows that detection of neoplasm in the blood is a path to early detection of cancer. While this could in principle be done using SNVs from panels of driver genes^35^ or with deep whole genome sequencing of cell free DNA^36^, the first will miss many cases and the second would be prohibitively expensive.

To carry out a MSLV assay, biological templates from samples would be tagged, enriched for selected loci with panels, subjected to mutagenesis, replicated ideally with independently tagged first copies, and then amplified for sequencing. In such assays, one would add synthetic control templates with known microsatellite lengths to provide a measure of error rates, so that the limits of detection could be modeled, perhaps as we showed in this paper. We can identify some uncertainties in this plan. (1) Is there sufficient MSLV in tumors? (2) Is there too much instability in a tumor to get a clear read-out? (3) Is there too much somatic variation in blood for tumor signal to be seen?

We can also anticipate some answers. First, while there are hundreds of thousands of microsatellites in the genome, assays based on only a few thousand might easily suffice for cancer detection. We estimate from the literature, and from our own internal tumor sequence data, that at least 1% of mononucleotide tracts have length variation in primary cancers^3-6^, much higher of course in patients with MS instability (MSI) syndromes^1,2^. We think it highly likely that this is an underestimate, as we have found in preliminary data that many of the mononucleotide C tracks, especially the longer ones, cannot be correctly read or even covered by standard sequencing libraries and platforms. Moreover, panels can be adjusted to increase the representation of highly variable loci. Even if the variation were only 1% sufficient numbers of variant markers would be present after enrichment of a few thousand loci. Second, if the mononucleotide tracks are too variable in a given cancer, for example in patients with high levels of MSI, the panels can be designed to include more stable microsatellites, such as dinucleotide tracts, or shorter tracts of mononucleotides. Third, while we expect to see some somatic variation in cell-free DNA in healthy individuals, we expect those variant length profiles will be relatively stable over time and distinct from the emerging new variants of a neoplasm.

## Data Availability

Sequence data is available in the Sequence Read Archive, BioProject accession number PRJNA763883. Processed data tables (read, well-covered first copy, and template tables) are available in Zenodo, https://doi.org/10.5281/zenodo.5149074. Code is available as a Supplementary Data file.

## Acknowledgements

We thank Siran Li for providing details on bisulfite mutagenesis, and Peter Andrews for providing analysis of microsatellites in the reference genome. We also thank Antoine Gruet and Inessa Hakker for their assistance generating sequencing libraries.

