## Supplementary Figures for "Accurate measurement of microsatellite length by disrupting its tandem repeat structure"

### Supplementary Figures and Tables

**Supplementary Figure 1: Reads per first copies.**

For each of the 5 libraries, we show the reads per first copy distributions used to establish a cutoff for minimum reads for a first copy to be well-covered. Each first copy has a point on the graph, with the x-axis indicating its position in a sorted list by number of reads, and the y-axis indicating the number of reads assigned to that first copy.

**Supplementary Figure 2: Sequencing base quality and base composition.**

For each of the 5 libraries, we split the raw sequencing reads into 4 sets, based on read number 1 or 2, and base composition of the microsatellite (A vs. T, C vs. G, CA vs. TG). For each set of reads, we show the per base Phred-scale quality score, as well as the per base composition of A/C/G/T. In the base quality plot, the blue line indicates mean base quality; the red line shows median base quality; the box plot shows the $25^{th}$ and $75^{th}$ quantiles in yellow and the $10^{th}$ and $90^{th}$ quantiles in black. The microsatellite region is highlighted in blue in both plots. Base quality drops dramatically after reading through the mononucleotide microsatellites. This impact on base quality is not observed in the mutated libraries.

**Supplementary Figure 3: Drop-out and matching tract lengths.**

For each of the unmutated mononucleotide tracts, M-17 (A-) and M-18 (C-), we show a scatterplot of the measured lengths from the two reads in the pair, with the length measured from the read reading the mono tract as A or C track on the x-axis and the length measured from the read reading the mono tract as a T or G track on the y-axis. The -1 position is reserved for those cases where both strands were unable to make a microsatellite call. This plot contains a down-sampling to 1 million data points.

**Supplementary Table 1: Sequence information of the templates and the oligonucleotides**

The sequences for the 3 template sequences: M-18 (C), D-26 (CA), and M-17 (A) are listed. N's indicate random nucleotides; D's indicate random nucleotides excluding C. Oligo (1) and primer sequences UP1 and UP3 are also listed for the two protocols used.

**Supplementary Table 2: Disruption yields as a function of mutation rate**

For the M-18 (C+) and D-26 (CA+) templates, the proportion of reads passing each of the two disruption parameter thresholds is modeled as a function of overall mutation rate. The reads that pass the rate cutoff are the reads with between 15% to 85% of C's mutated, and the reads that pass the repeat length cutoff have 5 or fewer units of the repeat intact. Highlighted in green are the mutation rate bins that most closely match the observed data.

**Supplementary Table 3: Companion table to Figure 3, per template first copy on- and off-target counts**

The data underlying Figure 3, for the libraries M-18 (C++), D-26 (CA-), and D-26 (CA++), is shown. Each template is tabulated by the number of well-covered first copies on-target (equal to 18 or 26) and the number of well-covered first copies off-target. For the disrupted libraries, templates have either all first copies on-target or all first copies off-target. For the D-26 (CA-), many templates are mixed with both on-target and off-target first copies.

**Supplementary Table 4: Lengths of unanimous templates**

For the three libraries, M-18 (C++), D-26 (CA-), and D-26 (CA++), shown in Figure 3, the number of templates with unanimous first copies, for each possible length, is tabulated, separated further by the number of first copies per template. The vast majority of unanimous templates are of the expected on-target length (18 or 26). Synthetic variant templates (3+ unanimous first copies) were removed before the final error rate estimations.

**Supplementary Table 5: Error rates as a function of disruption parameters**

For the two mutated libraries, we analyze the read off-target rate, and number of reads retained, as a function of the two disruption parameters. The max repeat length is varied from 3 to 18 for the M-18 library and 3 to 13 for the D-26 library, with the conversion rate thresholds varying from 0.45 to 0.55 as the strictest threshold, to 0.05 to 0.95 to the loosest threshold. Highlighted in green are the values for the parameter range in our main analysis. Error rates can be reduced further by using stricter disruption thresholds, though yield is also affected by this choice.

**Supplementary Table 6: Error rates from paired first copy observations, averaged rates**

For the four libraries, M-18 (C-), M-18 (C++), D-26 (CA-), and D-26 (CA++), we counted all paired first copy observations (L, K) where L is the MS length from one read, K the length from the other. To create an emulation in which all templates are weighted equally from the entire data set, we normalized counts per template, that is, for every template we determined the number of paired counts that were (L, K) and divided by the total. This generates a matrix over all possible (L, K) pairs for every template. These distributions are then averaged and shown for each library in this table.

**Supplementary Table 7: Error rates from paired first copy observations, emulated counts**

For the four libraries, M-18 (C-), M-18 (C++), D-26 (CA-), and D-26 (CA++), we counted all paired first copy observations (L, K) where L is the MS length from one read, K the length from the other. To create an emulation in which all templates are weighted equally from the entire data set, we normalized counts per template, that is, for every template we determined the number of paired counts that were (L, K) and divided by the total. This generates a matrix over all possible (L, K) pairs for every template. These distributions are then averaged. To emulate low-coverage paired first copy observations over N templates, we multiply this table by N, and round to the nearest integer, as shown for N=10,000 in this table.

Supplementary Figure 1

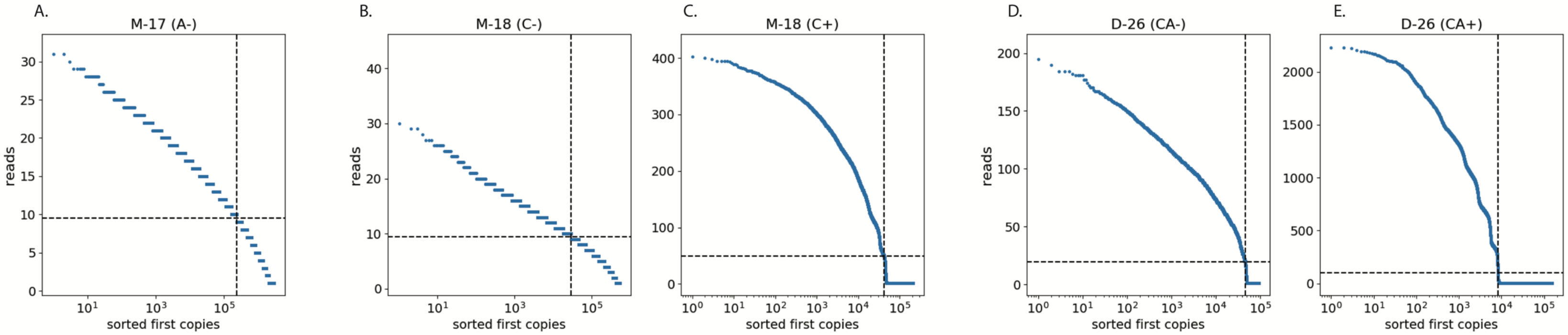


**Supplementary Figure 1: Reads per first copies.** For each of the 5 libraries, we show the reads per first copy distributions used to establish a cutoff for minimum reads for a first copy to be well-covered. Each first copy has a point on the graph, with the x-axis indicating its position in a sorted list by number of reads, and the y-axis indicating the number of reads assigned to that first copy.

### Supplementary Figure 2

#### Sequencing Base Quality and Base Composition

##### Annotated Example:

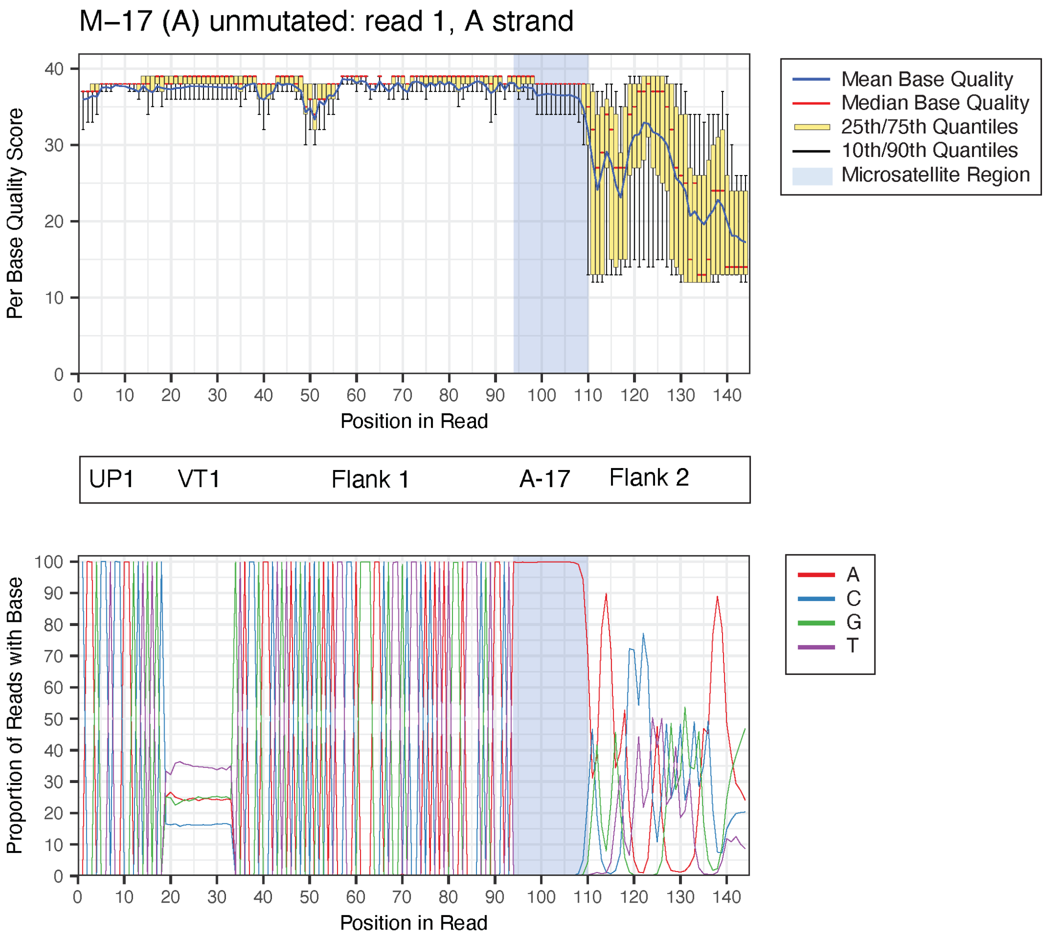


**Supplementary Figure 2: Sequencing base quality and base composition.** For each of the 5 libraries, we split the raw sequencing reads into 4 sets, based on read number 1 or 2, and base composition of the microsatellite (A vs. T, C vs. G, CA vs. TG). For each set of reads, we show the per base Phred-scale quality score, as well as the per base composition of A/C/G/T. In the base quality plot, the blue line indicates mean base quality; the red line shows median base quality; the box plot shows the 25$^{th}$ and 75$^{th}$ quantiles in yellow and the 10$^{th}$ and 90$^{th}$ quantiles in black. The microsatellite region is highlighted in blue in both plots. Base quality drops dramatically after reading through the mononucleotide microsatellites. This impact on base quality is not observed in the mutated libraries.

**A.**

### M-17 (A-) unmutated

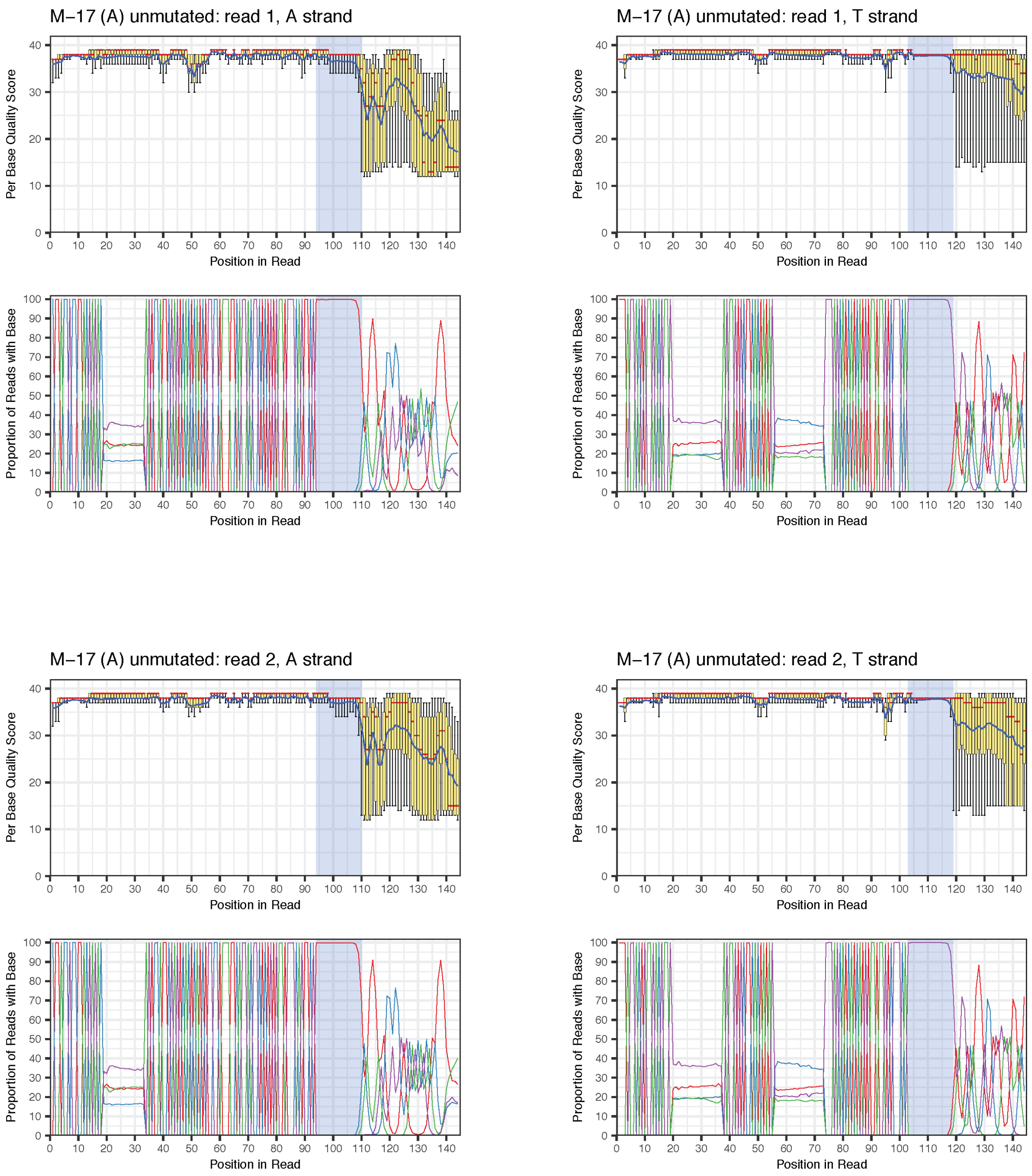

**B.**

### M-18 (C-) unmutated

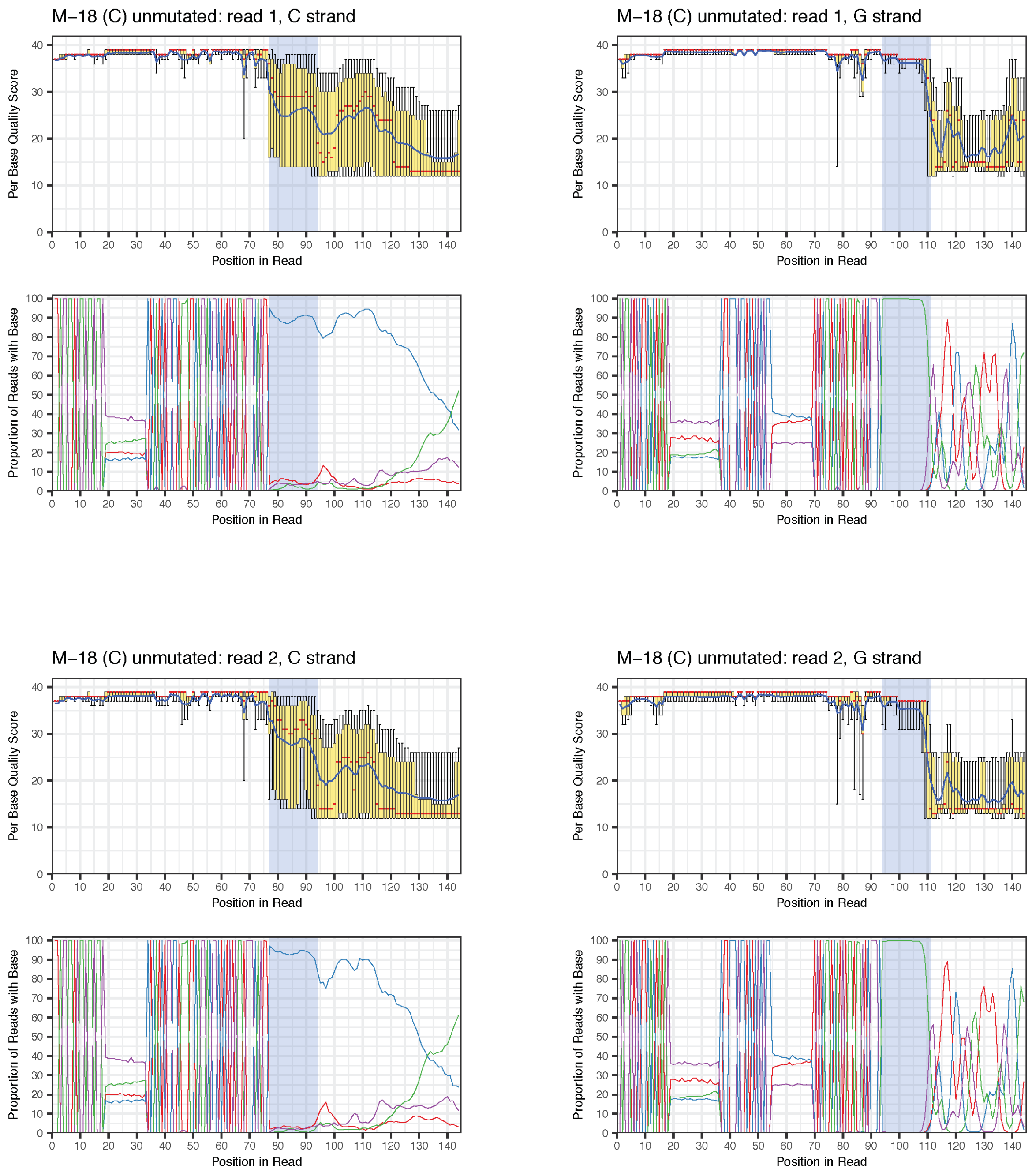

**C.**

### D–26 (CA-) unmutated

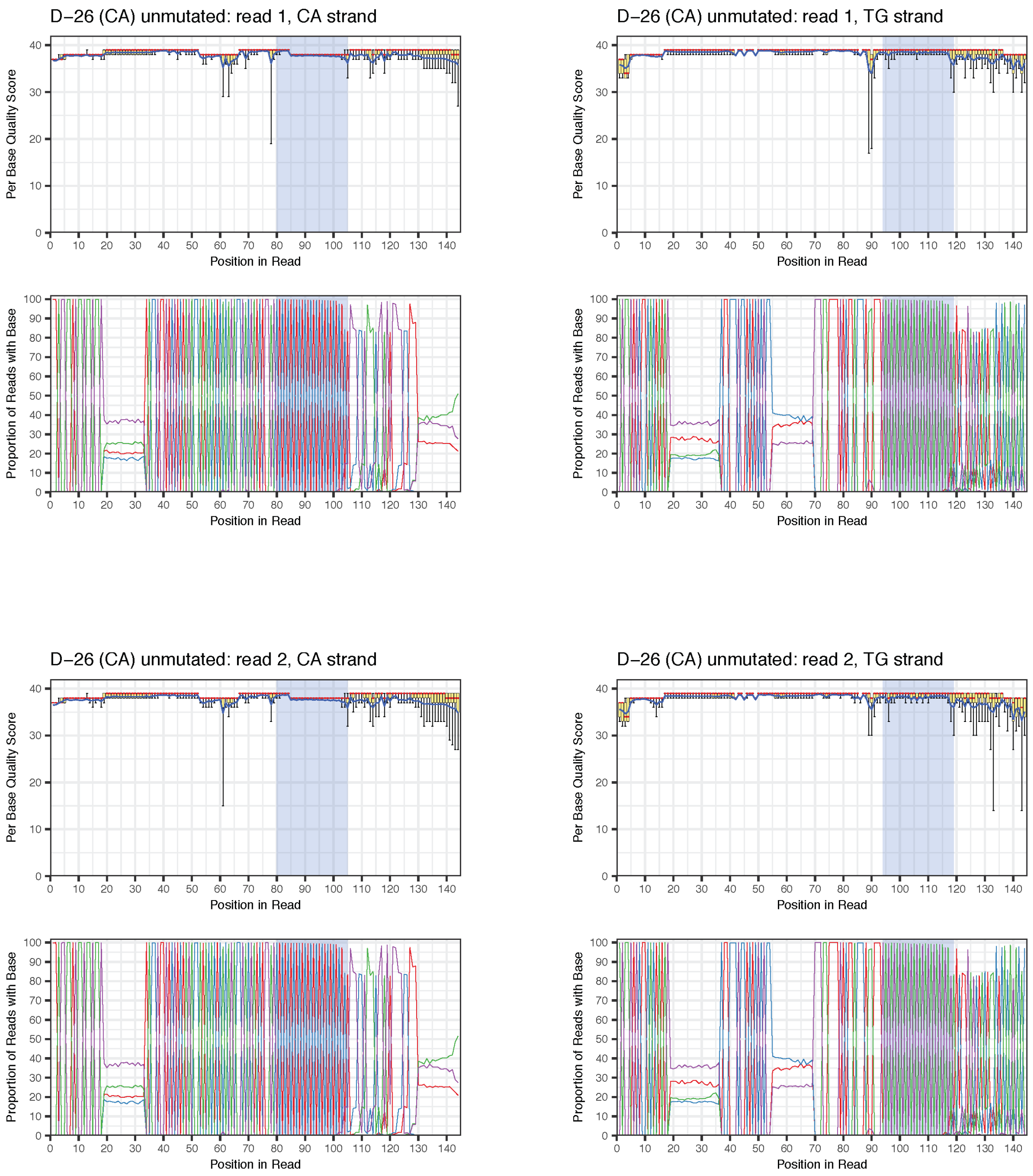

**D.**

### M-18 (C+) mutated

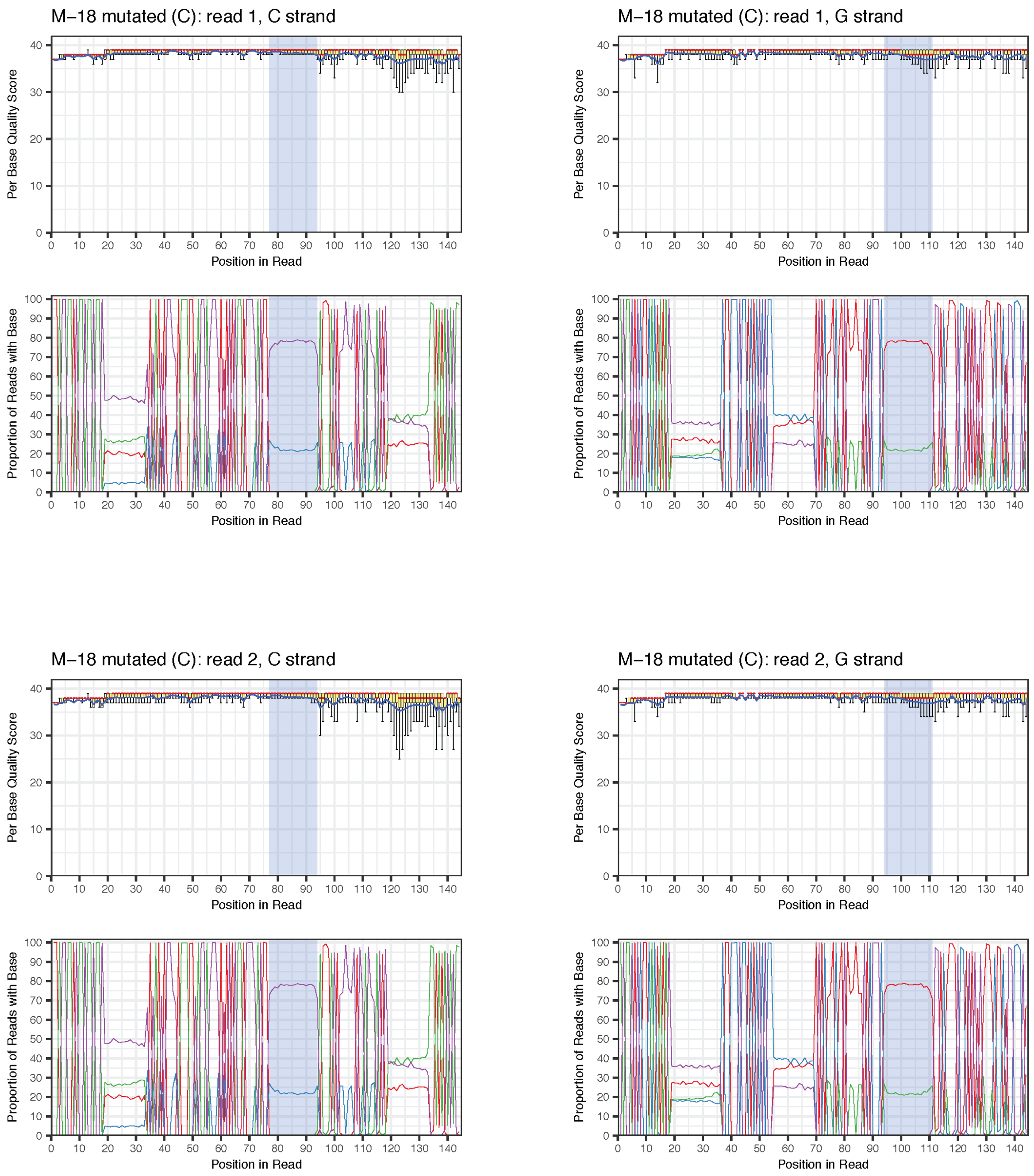

E.

### D–26 (CA+) mutated

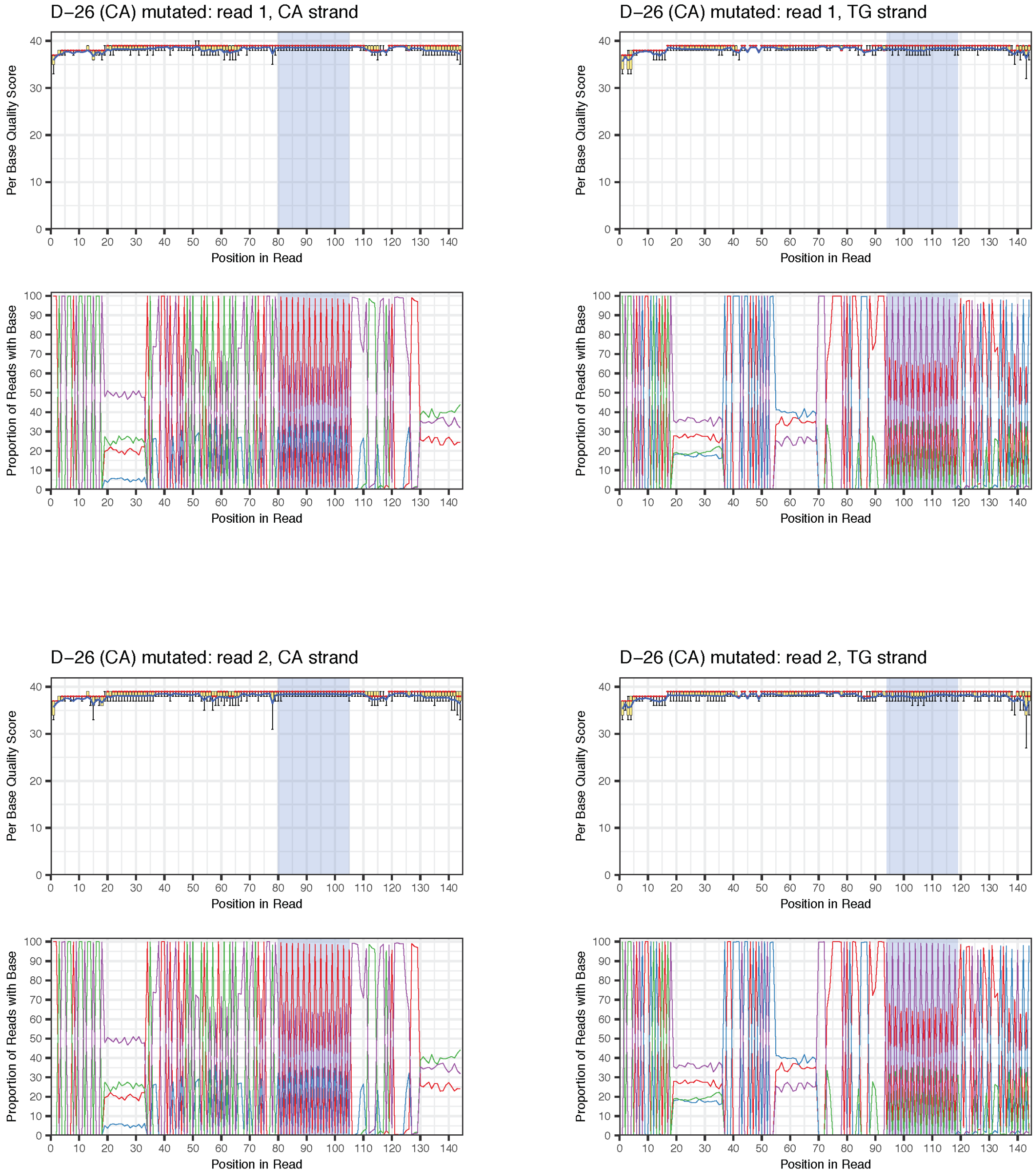

### Supplementary Figure 3

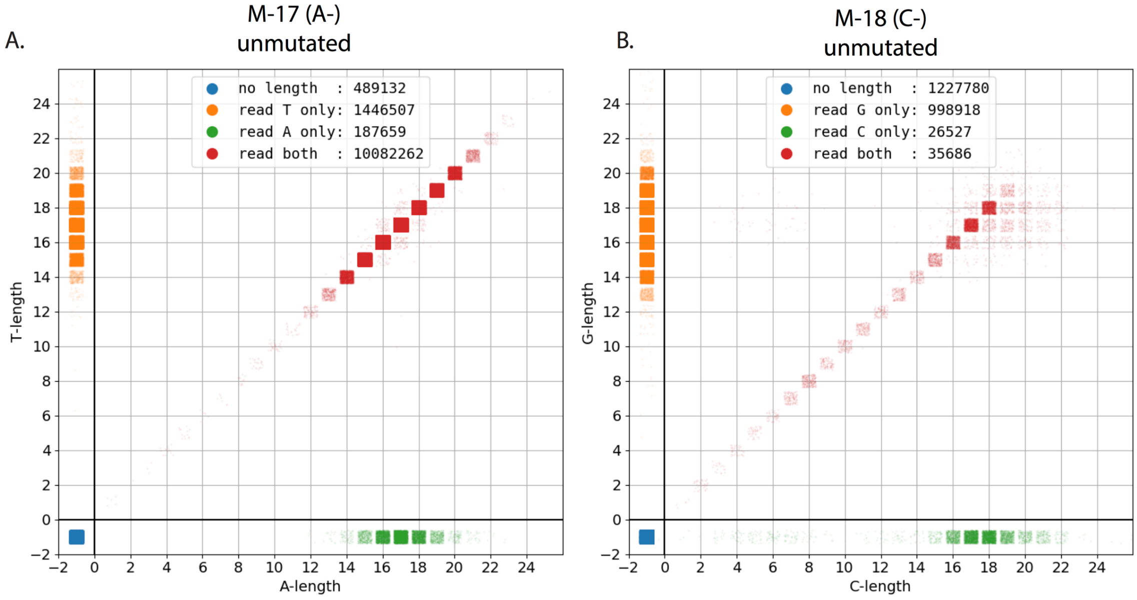


**Supplementary Figure 3: Drop-out and matching tract lengths.** For each of the unmutated mononucleotide tracts, M-17 (A-) and M-18 (C-), we show a scatterplot of the measured lengths from the two reads in the pair, with the length measured from the read reading the mono tract as A or C track on the x-axis and the length measured from the read reading the mono tract as a T or G track on the y-axis. The -1 position is reserved for those cases where both strands were unable to make a microsatellite call. This plot contains a down-sampling to 1 million data points.
