## Supplementary Data for "Accurate measurement of microsatellite length by disrupting its tandem repeat structure"

```
## Luria-Delbruck Diffusion Simulation Code

import numpy as np
import matplotlib.pyplot as plt
import matplotlib
import time

default_color_list = matplotlib.rcParams['axes.prop_cycle'].by_key()['color']

np.random.seed(hash( "This is not a random seed." ) % 2**32)

def simulate_LD_diffusion(N, R, copy_rate, error_rate, error_up,
FIRST_ROUND_ERROR=False, VERBOSE=False):
    if FIRST_ROUND_ERROR:
          ## if there's first round error, we start with 3 columns (including
      one up and down)
        C = 3
        ## and we seed X with the results of a multinomial draw with the
        ## given error rate (assume first copy happened)
        has_error      = np.random.random(N) < error_rate
        has_up_error   = np.random.random(N) < error_up
        X = np.zeros(shape=(N, C), dtype=int)
        X[has_error*(~has_up_error), 0] = 1
        X[~has_error                , 1] = 1
        X[has_error*has_up_error    , 2] = 1
        MSLV = [-1, 0, 1]
    else:
        ## number of columns, initially 1
        C = 1
        MSLV = [0]
        ## X will grow with each round to record the count per length.
        ## we start with one copy at the starting length for each simulation.
        X = np.ones(shape=(N, C), dtype=int)
    for r in range(R):
        ## first, how many copies at this spot
        num_copies = np.random.binomial(X, copy_rate)
        ## of those copies, how many are error
        num_copies_error = np.random.binomial(num_copies, error_rate)
        ## of the errors how many are up
        num_copies_up    = np.random.binomial(num_copies_error, error_up)
        ## the rest of the errors are down
        num_copies_down  = num_copies_error - num_copies_up
        ## copies without errors stay here
        num_copies_equal = num_copies - num_copies_error
        ## prepare the next matrix with two extra columns
        ## expanding down one (-1) and up one (+1)
        Y = np.zeros(shape=(N, C+2), dtype=int)
```

```
        ## down copies added to the left of the matrix
        Y[:,  :-2] += num_copies_down
        ## equal copies and the previous matrix added to the center
        Y[:, 1:-1] += num_copies_equal + X
        ## up copies added to the right of the matrix
        Y[:, 2:  ] += num_copies_up
        ## check if we need to have Y get wider:
        ## check the upper and lower bounds of Y to see if they are occupied
        Ysum = np.sum(Y, axis=0)
        ## if not, trim
        if Ysum[0] == 0:
            Y = Y[:,1: ]
        else:
            ## if yes, then keep add it to the bounds and increment C
            MSLV.insert(0, MSLV[0]-1)
            C += 1
        ## check the upper end, if there's nothing new, don't extend
        if Ysum[-1] == 0:
            Y = Y[:   , :-1]
        else:
            MSLV.append(MSLV[-1] + 1)
            C += 1
        ## replace X with Y
        X = Y
        if VERBOSE:
            print r,
    if VERBOSE:
        print
    return X, MSLV

FIRST_ROUND_ERROR = True

N = 1000000
R = 22
copy_rate  = 0.95
eplus      = 0.001
eminus     = 0.01
error_rate = (eplus + eminus)
error_up = eplus / error_rate

t0 = time.time()
X, MSLV = simulate_LD_diffusion(N, R, copy_rate, error_rate, error_up,
      FIRST_ROUND_ERROR=FIRST_ROUND_ERROR, VERBOSE=True)
read_total =             np.sum(X, axis=0)
read_rate  =             read_total / float(np.sum(read_total))
best_call        =       np.argmax(X, axis=1)
first_copy_count =       np.bincount(best_call, minlength=(len(MSLV) + 1))
first_copy_rate  =       first_copy_count / float(N)
```

```
best_rate = np.max(X, axis=1) / np.sum(X, axis=1).astype(float)
MSLV = np.array(MSLV)

center = np.where( MSLV == 0 )[0][0]
bins = np.linspace(0, 1, 100)


fig = plt.figure(figsize=(14, 5))
for ind, bc in enumerate( [center-1, center, center + 1] ):
    ax = fig.add_subplot(1, 3, ind+1)
    bc_filter = best_call == bc
    label = " MSL = %d; %0.4f" % (MSLV[bc], np.sum(bc_filter) / float(N))
    ax.hist(best_rate[bc_filter], bins=bins, label=label, density=True ,
      alpha=1, color=default_color_list[ind])
    plt.legend(fontsize=15)

plt.show()
```
